# *In-vivo* monitoring and quantification of breast cancer growth dynamics with non-invasive intravital mesoscopic fluorescence molecular tomography

**DOI:** 10.1101/2020.08.03.234898

**Authors:** Mehmet S. Ozturk, Marta G. Montero, Ling Wang, Lucas M. Chaible, Martin Jechlinger, Robert Prevedel

## Abstract

Preclinical breast tumor models are an invaluable tool to systematically study tumor progression and treatment response, yet methods to non-invasively monitor the involved molecular and mechanistic properties under physiologically relevant conditions are limited. Here we present an intravital mesoscopic fluorescence molecular tomography (henceforth IFT) approach that is capable of tracking fluorescently labeled tumor cells in a quantitative manner inside the mammary gland of living mice. Our mesoscopic approach is entirely non-invasive and thus permits prolonged observational periods of several months. The relatively high sensitivity and spatial resolution further enable inferring the overall number of oncogene-expressing tumor cells as well as their tumor volume over the entire cycle from early tumor growth to residual disease following the treatment phase. We find that sheer tumor volume, as commonly assessed by other imaging modalities, is not well correlated to tumor cell quantity, hence our IFT approach is a promising new method for studying tumor growth dynamics in a quantitative and longitudinal fashion *in-vivo*.

## Introduction

The current frontiers in the battle against breast cancer are shifted towards early detection on the one hand and understanding mechanisms that lead to recurrent tumors on the other hand. While therapeutic improvements in combination with early tumor detection enabled through the introduction of mass mammographic screening programs has led to a substantial increase in breast cancer survivors over the last few decades^1^, a detailed, longitudinal insight into early tumor initiation events - like ductal carcinoma in situ (DCIS) - under physiological conditions is still lacking^2^. A further major problem in modern breast cancer care is constituted by the increase of breast cancer related death due to recurrent disease^3^. These incurable recurrences are caused by treatment refractory cells that stay dormant over years^4,5^. Similar to the situation at the tumor induction phase, these small amounts of residual cells are hard to trace and molecular markers of this population need to be established to understand potential points of detection and therapeutic interference^6^. Such information may be difficult to decipher solely by studying cancer in the human population, due to the impossibility to follow scarce cells in these cancer stages for mechanistic analyses in an enormously heterogeneous patient population.

To this end, preclinical mouse models can provide valuable resources for cancer prevention and treatment strategies, since they enable both longitudinal and molecular analyses of precancerous and cancerous lesions in defined genetic contexts and in environmentally-controlled conditions, which would be exceedingly difficult to accomplish solely by studying human cancer^7,8^. Preclinical breast cancer mouse models, such as transplantation based mouse models or genetically engineered mouse models (GEMMs) have been successfully used to investigate tumor biology via a range of imaging approaches^9^ and therapeutic strategies^10^. Moreover, GEMMs based on conditional gene targeting wherein genes of interest are inactivated (or activated) in a temporal and tissue-specific manner offer advanced tools to study cancer with the inclusion of reporter alleles to perform *in-vivo* imaging^11^. Finally, recent advances in mammary tumor biology with respect to transplantation^12^ and intraductal injection techniques^13^ permit the study of cells derived from above mentioned models as well as genetically engineered lines (**Fig. 1a,b**).

**Fig. 1.**
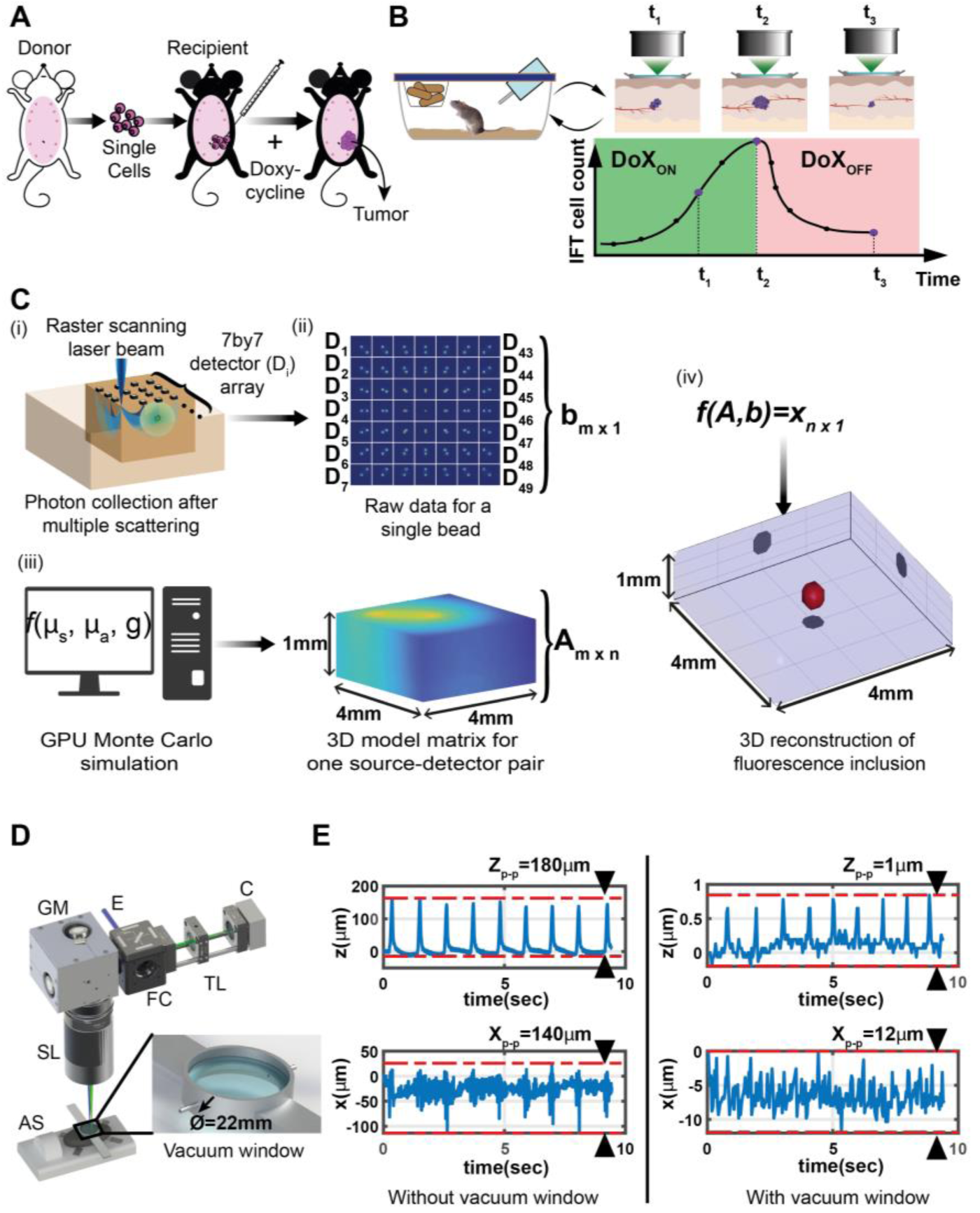
Experimental outline and schematic representation of the principles of the Intravital Fluorescence Tomography (IFT) system. A. Extraction of tumor cells from the donor animal and the implantation into the receiving animal. B. Schematic of longitudinal imaging of a single animal. Different time points refer to different stages of tumor development: tumor growth (Doxycycline administration: DoXON) and tumor regression (DoXOFF) C. Principle of IFT method: (i) Focused laser beam excites fluorescence inside the sample volume by raster scanning while a detector array simultaneously images the scanned area. For each scan location, multiple-scattered emission light is collected by the detector array (sCMOS camera). (ii) After a full scan of the sample, each detector yields an image, with all detector images together constituting the raw data (measurement). (iii) A GPU-accelerated Monte Carlo algorithm simulates the optical propagation of scattered light inside the sample volume. (iv) The raw data and model matrix are fed into a reconstruction algorithm, to extract the 3D distribution of the fluorescence emission. D. A CAD representation of the IFT imaging setup. An excitation laser (E) enters the system through a standard filter cube (FC). The laser is raster scanned by a 2D galvanometric mirror (GM) and imaged onto the sample by a scanning lens (SL). The emitted fluorescence light is de-scanned, separated from the excitation light by the FC and imaged onto the camera (C) by an imaging lens (IL). For *in-vivo* experiments we use an anesthesia system (AS) for narcosis and monitoring of physiological parameters while the region of interest is stabilized with a custom, non-invasive imaging window (NIVOW – zoom-in). E. Mouse skin position as a function of time as measured by OCT, showing tissue motion along z and x axes without (left) and with (right) NIVOW.

This extensive collection of available pre-clinical breast cancer models represents a powerful toolset to explore mechanisms on tumor evolution and therapy success. Despite their unique advantages, current studies employing preclinical mouse models do not harness their full potential. This is because currently employed methods to visualize and assess tumor progression have important limitations with regards to either sensitivity, resolution, imaging field-of-view (FOV) and depth or the ability to monitor disease over prolonged timeframes. In particular, histological approaches, while providing valuable quantitative, high-resolution information on tumor morphology and abundance, are end-point methods and thus prohibit reconstruction of individual, longitudinal tumor development. This ability, though, is critical in assessing tumor populations during early growth phases as well as during treatment to understand dynamics of shrinkage and evolution of resistant disease. Intravital, longitudinal imaging techniques, on the other hand, can give valuable information on the temporal evolution and dynamics of cell populations^14^. But because of the highly scattering nature of the mouse skin and tumor tissues, optical access for standard light microscopy methods such as confocal or two-photon has to be guaranteed by either surgical implantation of so-called imaging windows^15,16^ or by applying windows on surgically exposed organs^17–21^. In addition to the limitations in imaging FOV and depth, these surgical procedures, however, typically trigger inflammatory responses that may alter the tumor micro-environment^15^, are limited in their respective size and bear significant burden for the animal when utilized over extended time-frames. Furthermore, as these windows are static, they are not fully compatible with spontaneous mouse breast tumor models, as these cases require dynamic and flexible repositioning of the imaging field-of-view (FOV) in order to monitor stochastic tumor emergence over ten mammary glands.

An alternative method for non-invasive monitoring of tumor burden and screening for drug efficacy in cancer mouse models is bioluminescence imaging (BLI). Although its non-invasive character and high sensitivity are promising features^22^, the spatial resolution achievable in 3D reconstructions is typically very poor (∼mm’s)^23^. Furthermore, the bioluminescent signal mainly depends on two components: the substrate/enzyme amounts and their interaction, as well as the accessibility of oxygen in the microenvironment. Both parameters are variable and challenging to assess independently, therefore the normalization of bioluminescent signal required for extracting quantitative information has remained challenging. While quantification attempts were shown in 2D imaging^24^, tomographic studies are predominantly qualitative^25^ to date, and have instead capitalized on the high-throughput nature and sensitivity of the method. Furthermore, BLI requires injections for every imaging experiment in a longitudinal time-series, and thus puts additional burdens on the animal.

Besides the above-mentioned fluorescent and luminescent based techniques, other mature technologies are also being used for monitoring tumor development. These conventional techniques include computed tomography (CT), magnetic resonance imaging (MRI), and ultrasound (US) and typically only deliver unspecific structural information with limited spatial resolution^26^. Despite recent advancement that enable cellular specificity through the use of engineered molecular constructs^27^, the achievable spatial resolution is again limited to a few hundred microns. Furthermore, for CT and MRI, achieving high spatial resolution requires prolonged imaging sessions that may last over several hours, which puts additional burden on experimental animals. Optical spectroscopy techniques can be regarded as an alternative non-invasive, fast and low-cost approach^28^. As these methods quantify the spatiotemporal changes of physiological parameters (e.g. oxy-/deoxy-hemoglobin concentration, oxygen saturation etc.) of the tumor environment^29^, they however only provide surrogate (e.g. metabolic) information about the state of tumor cells. Another recently emerging technique is photoacoustic imaging (PAI) which is capable of detecting absorbing contrast agents deep (∼1cm) in scattering tissue with mesoscopic resolution (a few 100µm). PAI is inherently sensitive to chromophore contrast so it is suitable for imaging vascularization and their oxygenation state via intrinsic spectral absorbance differences of (oxy-/deoxy-) hemoglobin. Moreover, the recent introduction of gene reporters for molecular imaging in Photoacoustic Tomography (PAT) offers the possibility of non-invasive imaging with cellular specificity and high sensitivity (∼100-1000 cells)^30,31^. Although multispectral PAT is rapidly being developed, current PAT implementations involve costly and custom-made components and its operation entails complicated post-processing, thus limiting its use to a handful of labs worldwide.

In this work, we present an experimental approach that aims to address many of the above-mentioned shortcomings. Our platform, named IFT, utilizes an optical tomography approach known as mesoscopic fluorescence molecular tomography (MFMT)^32–38^, and includes a number of key improvements that enable quantitative and longitudinal studies of *in-vivo* tumor development in inducible mouse models. First, inspired by Looney et al. ^18^, we introduce a modified, non-invasive imaging window that minimizes motion artifacts induced by breathing of the animal and ensures repeated imaging of the same location throughout the study with high stability. Second, subtle changes to the opto-electronic design of our platform compared to previous implementations^39,40^ enables larger scan areas (FOVs) at twice improved data acquisition rates. Third, after careful characterizing our IFT tomographic resolution and sensitivity, we calibrate its signal with both *in-vitro* phantoms and histological tumor samples, thereby establishing a linear relationship from which we can infer the number of fluorescently labelled, i.e. tumorigenic, cells with high (∼200µm) mesoscopic resolution in 3D. Although previous works have monitored tumor xenografts at single timepoints using MFMT^35,40–42^, here for the first time, we report longitudinal imaging of the entire tumor life cycle spanning several weeks *in-vivo*.

We show the capability of IFT to monitor tumor progression over several months in a non-invasive, longitudinal fashion by applying it to a preclinical mouse model of inducible breast cancer. In particular, we track tumor cell number as well as tumor volume and geometry through an entire cycle of tumor growth and tumor regression upon silencing of the oncogenic signal. Here, the high sensitivity of our technique enables tumor identification long before they become palpable, thus making it possible to investigate aspects of early tumor growth and also treatment effects monitoring survival of refractory subpopulations, while the high 3D spatial resolution enables assessing subtle changes in tumor geometry over time. Most remarkably, however, we find that overall tumor volume, as commonly measured by other cancer imaging modalities, does not correlate well with the tumor cell quantity, hence questioning the ability of these methods to accurately assess tumor burden. Therefore, we believe IFT to be a promising new method for studying breast tumor biology in a quantitative and longitudinal fashion *in-vivo*.

## Results

### Principles of intravital fluorescence tomography

Intravital Mesoscopic Fluorescence Molecular Tomography (i.e. intravital MFMT, or henceforth in short IFT) excites fluorescence via one-photon absorption and collects multiple scattered photons on a 2D detector array (**Fig. 1c**). The data collection principle is conceptually similar to previous implementations of MFMT ^39,43^. An excitation laser beam (561nm) is focused on the sample and raster-scans an excitation spot across the sample surface. For each scan point, a camera records a wide-field image of the sample, containing all the backscattered/fluorescent photons. Each camera pixels (detector) collects photons at a varying distance away from the laser (source) (**Fig**.**1c, i**). With increasing detector distance, the collected photons predominantly originate from increasing depths in the sample, thus the resulting camera image carries intrinsic depth (axial) information. In order to reconstruct a meaningful 3D image from the raw 2D data sets, IFT is based on a so-called forward model (Jacobian matrix), which is calculated by simulation of photon propagation inside the scattering tissue of interest via a GPU-accelerated Monte-Carlo method^37,44^ (**Fig. 1b,iii** and **Methods**). The optical properties of mammary gland tissue and skin were measured by a home-built dual integrating-sphere setup^45^ (**Supplementary Note 1 and Supplementary Fig. S1 & S2**), and the so obtained values were used as a starting point for the calculation of the Jacobian matrix, which was further optimized to ensured accurate and reliable 3D reconstructions. The Jacobian was then used to retrieve the 3D distribution of the fluorescence emitters from the experimental imaging dataset by running an Lp-norm reconstruction algorithm^46,47^ **(Fig. 1c,iv** and **Methods**).

### Non-invasive imaging window

Intravital imaging approaches typically involve the surgical implantation of an imaging windows which can trigger inflammatory responses and bear significant burden for the animal when utilized over extended time-frames^15–21^. Moreover, one recurring challenge in intravital imaging is breathing-induced tissue motion, which commonly leads to severe artefacts when not properly accounted for. While the image acquisition can in principle be synchronized or gated to the periodic movement of the body cavity^48^, such techniques are cumbersome and require specialized equipment, are prone to error and typically result in prolonged imaging times as the tissue is at rest only during a small fraction of the breathing cycles. To overcome these challenges, we designed a non-invasive, vacuum-operated stabilization window (NIVOW), inspired by^18^. It comprises a round metal ring (22mm inner diameter) sealed with a coverslip that can be placed on the mouse skin at the center of the imaging FOV (**Fig. 1d**). Vacuum can be applied through two small openings, which firmly attaches the skin to the coverslip, while the window housing is secured to the optical table via suitable arms. This approach allowed us to leave the mouse skin intact (i.e. no surgical intervention) and enabled repeated and flexible positioning for imaging. Furthermore, the vacuum stabilizes the region of interest and its underlying tissue significantly, both laterally as well as axially (**Fig. 1e**). Quantitatively, the axial motion of skin and subcutaneous tissue layers decreased from over >100µm to ∼1µm peak-to-peak, as measured at high-speed by Optical Coherence Tomography (OCT) (**Supplementary Note 2** and **Supplementary Video 1 & 2**). This residual motion is negligible compared to our spatial resolution and the coverslip further ensured a ‘flat boundary’ condition as required by the Jacobian model matrix. Thus, our NIVOW is an instrumental part of the IFT system for achieving high-fidelity reconstruction. Furthermore, it enables flexible repositioning of the imaging location. As the site of tumor onset can shift in the abdominal mammary gland area by several millimeters, having an adjustable region of interest brings a critical advantage over traditionally used, fixated imaging windows ^15^. Most importantly, however, no surgical intervention is required for placing the window and thus (i) no inflammatory signal can interfere with the tumor biology under investigation, and (ii) monitoring can be performed over the entire life-time of the animal.

### IFT characterization (resolution & sensitivity)

Our aim is to capture the process of breast cancer tumors *in-vivo* in its entirety: From the first formation of a solid tumor mass by a cluster of cells until the stage where the tumor becomes palpable (typically >5mm). For this our IFT imaging system must display the required spatial resolution as well as sensitivity. Initial tumor populations are commonly assumed to comprise ∼1×10^4^ cells, which hence marks our sensitivity requirement. Considering the average diameter of a mammary gland tumor cell (10-15µm), these smallest units of tumor populations approximates to 0.01 mm^3^ volume, hence necessitating a similar spatial resolution on the order of ∼250µm. In order to quantify the spatial resolution of IFT, we placed individual, 50µm fluorescent polymer microspheres (Cospheric, USA) under the mouse skin and imaged them with our IFT system (**Fig. 2a**). Reconstructing the acquired IFT image dataset indeed yielded single, bead like objects, which we used to characterize the point-spread function (PSF) of our system (**Fig. 2b**). In contrast to previous studies which often characterized spatial resolution by reconstructing a single focal object, here we purposely chose to reconstruct two objects in close proximity. Our characterization yielded a lateral (axial) resolution of ∼200-220µm (∼250µm), thus sufficient to resolve initial tumor clusters of ∼ 10,000 cells.

**Fig. 2.**
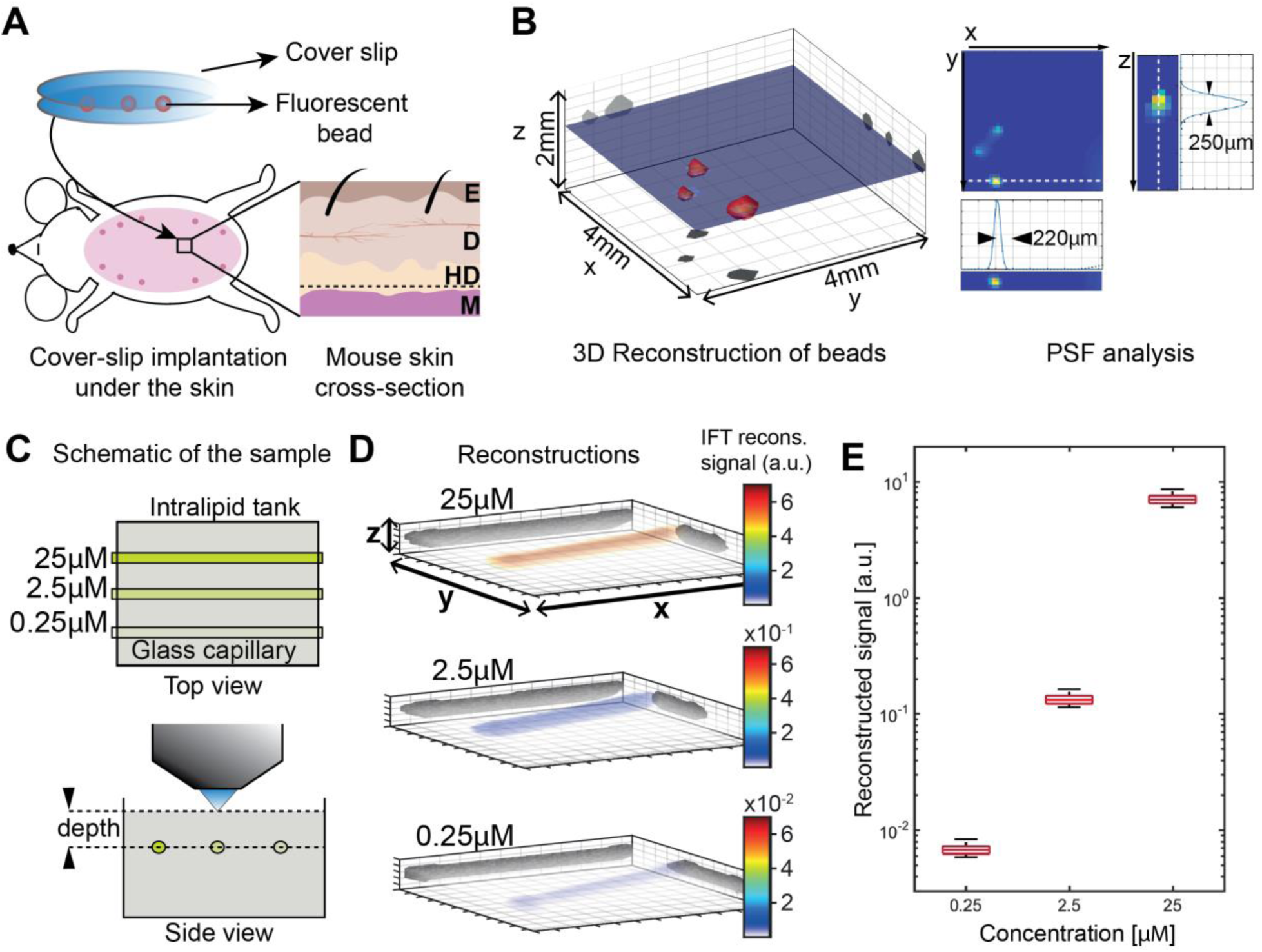
Characterization of IFT. A. Fluorescent beads, sandwiched between cover-slips, were placed under the mouse skin in the hypodermis (HD) under the epidermis ((E) and dermis (D), above the muscle (M). B. Spatial resolution analysis based on tomographic reconstruction of these beads. Point spread function (PSF) indicating lateral and axial resolution. C. *In-vitro* control experiment to establish linearity of the reconstruction algorithm. Glass capillaries filled with fluorescein of different concentrations (25, 2.5, and 0.25µM) inside an 10% Intralipid tank at 1.3mm depth were imaged by IFT. D. Reconstructions in 3D (x,y,z = 11×11×2mm3) for each concentration. Color bar indicates the reconstructed signal value (a.u.). E. The imaged concentrations show a linear relationship in their reconstructed IFT signal.

### Reconstruction algorithm linearity

One of the main aims of our study which goes beyond the capability of other established cancer imaging methods is to quantitatively measure the number of oncogene-bearing cells that contribute to the signal in IFT. A critical prerequisite for this is to show that the intensities of the IFT reconstructions scale linearly with the underlying fluorescence intensity. To investigate this relationship *in-vitro*, we engineered a scattering tissue phantom (Intralipid 10%) comprising fluorescein filled tubes (∼3µL) with three different concentrations (25, 2.5 and 0.25 µM) at a depth of 1.3mm (**Fig. 2c**). This depth was chosen since it is equivalent to most of the tumor location under the skin. IFT reconstructions (**Fig. 2d**) delivered signal levels that scaled linearly with actual fluorescein concentrations (**Fig. 2e**).

### Calibration of IFT cell count

Since in-vitro phantoms cannot replicate the real, *in-vivo* tissue heterogeneity, we next proceeded to calibrate our IFT signal values with absolute cell quantities obtained through tumor histology (**Fig. 3**). The presence of strong heterogeneity in nuclear H&E staining made it difficult to count nuclei with a standard cell counting software, therefore we devised a custom image processing pipeline (**Fig. 3a,b**). Here, we first restricted our analysis to a small sub-area of the tumor site that displayed homogenous H&E staining, and manually identified tumor clusters for automatic cell counting. This yielded crucial tumor cell density information (#cells/area). In parallel, the entire H&E slice was thresholded to separate tumor cells from other, stromal areas. Then, by measuring the respective entire tumor area and utilizing the cell density information, the overall number of cells pertaining to a given tumor section could be calculated. Likewise, this calculation was repeated for all histological sections, yielding an overall cell number for a 3D tumor. This pipeline was applied to (n=8) mice at different stages of tumor progression while mice were on DoX treatment. After IFT imaging of the tumor site, these mice were sacrificed, and tumor tissue extracted and prepared for the abovementioned histological Hematoxylin & Eosin (H&E) analysis. **Fig. 3c** shows histological (ground truth) number of cells plotted against the integrated IFT reconstructed signal for each tumor. We found the resulting graph to display a linear relationship with a fitting performance of R^2^=0.94 and RMSE=4.78, thus demonstrating the ability of our IFT system to faithfully capture the amount of fluorescent, i.e. oncogene bearing, tumor cells from the reconstructions. The fit to this data constituted our IFT calibration and was hence used in all experiments to convert reconstructed IFT signal to actual cell quantities. Furthermore, we note that the smallest tumor investigated during this calibration study was histologically measured to only contain ∼10^5^ cells. As we were also able to identify and reconstruct this small tumor mass via IFT this number also represents a conservative estimate of our *in-vivo* IFT sensitivity. Finally, IFT reconstructions and histological slices of the same tumor show considerable morphological overlap, further corroborating the capability of IFT to faithfully capture the 3D tumor geometry and volume (**Fig. 3c, right**).

**Fig. 3.**
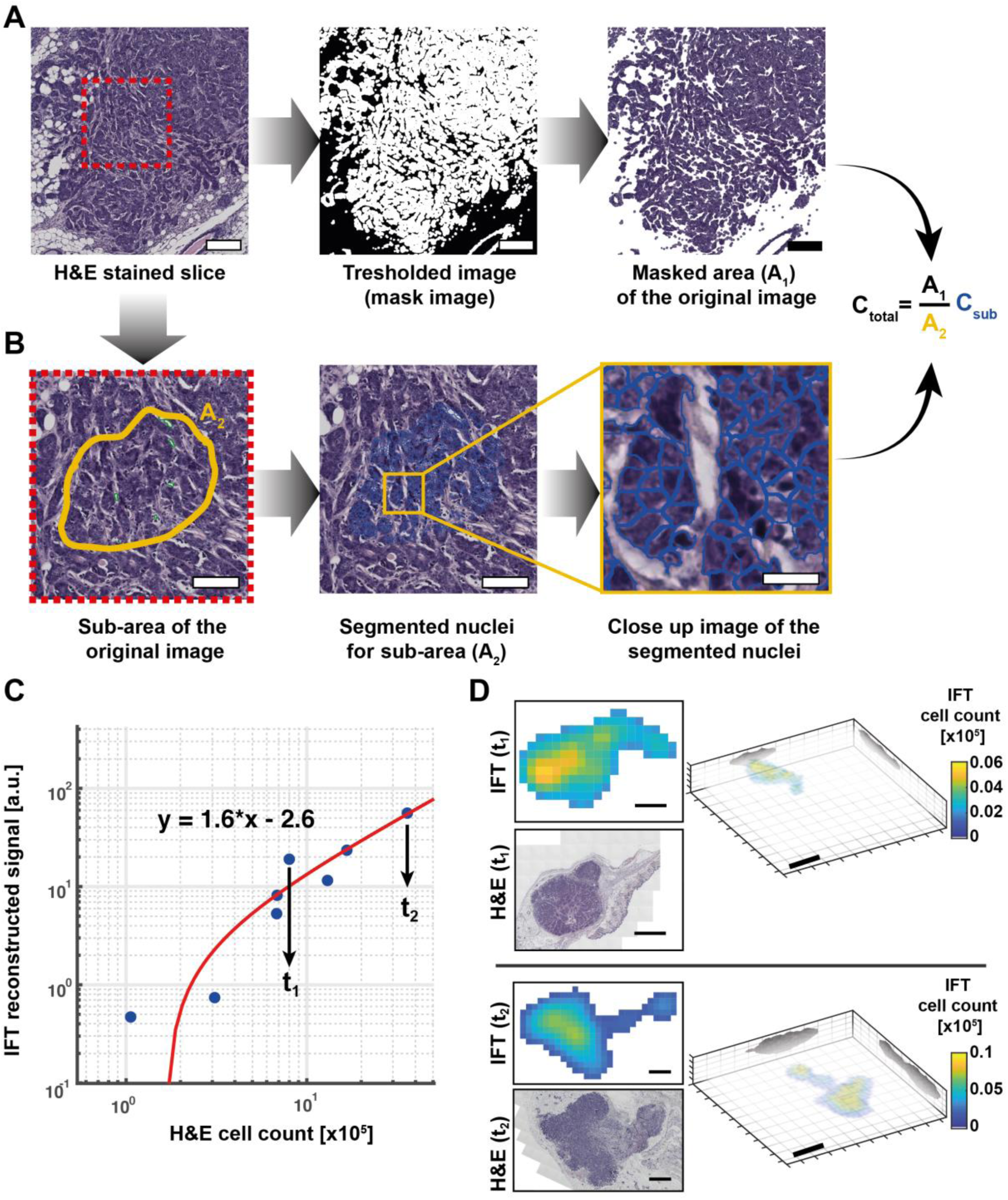
IFT cell quantification. A. Hematoxylin & Eosin (H&E) stained tumor slices were imaged and tumor coverage area (A1) was calculated via a threshold mask. Scale bar, 100µm. B. A subset of the image with good nuclear staining was chosen for automated cell counting, yielding tumor island area (A2) and cell count (Csub). Overall tumor cell number was then estimated as indicated. Scale bar, 50µm (zoom-in 20µm). C. Integrated IFT reconstruction signal values for 8 tumors of varying size and intensity plotted against ‘ground-truth’ cell count as obtained by H&E analysis. Linear fit to the data constitutes the IFT cell count calibration and was used for subsequent experiments. D. Exemplary 2D IFT cross-sections and corresponding H&E slices as well as 3D IFT reconstructions for two representative tumors in C. Scale bars 1mm (left) and 2mm (right, 3D reconstruction).

### Longitudinal molecular imaging of tumor development

After characterizing our IFT imaging system, we applied it to monitor mammary gland tumor development longitudinally (**Fig. 4**). For this, mice (n=2) with primary tumor cells implanted into their fat pad area were put on Doxycycline (DoX) for 4-5 weeks, throughout which the tumorigenic cells continued to grow (see timepoints t_1_-t_2_ in Fig. 4). This suspected growth phase was clearly recapitulated by a strong (∼10-fold) increase in IFT signal and hence tumorigenic cell quantity. Likewise, after the removal of DoX from the food diet, the cell numbers rapidly declined within a week (t_3_), very much similar to the dynamics observed in the original transgenic model^6^, and returned to or below base level within ∼40-50 days (t_4_). At this point (t_4_) the remaining, weak IFT signal can be attributed to scar tissue of the residual disease, which we therefore consider baseline (autofluorescence) signal. From these reconstructed data, we can also estimate the sensitivity of our system, which is on the order of 2-3×10^3^ cells per voxel. Remarkably, although we could observe a clear dependence of cell quantity on DoX, overall tumor volume did not show a clear correlation with the DoX cycle or cell quantity as assessed by IFT.

**Fig. 4.**
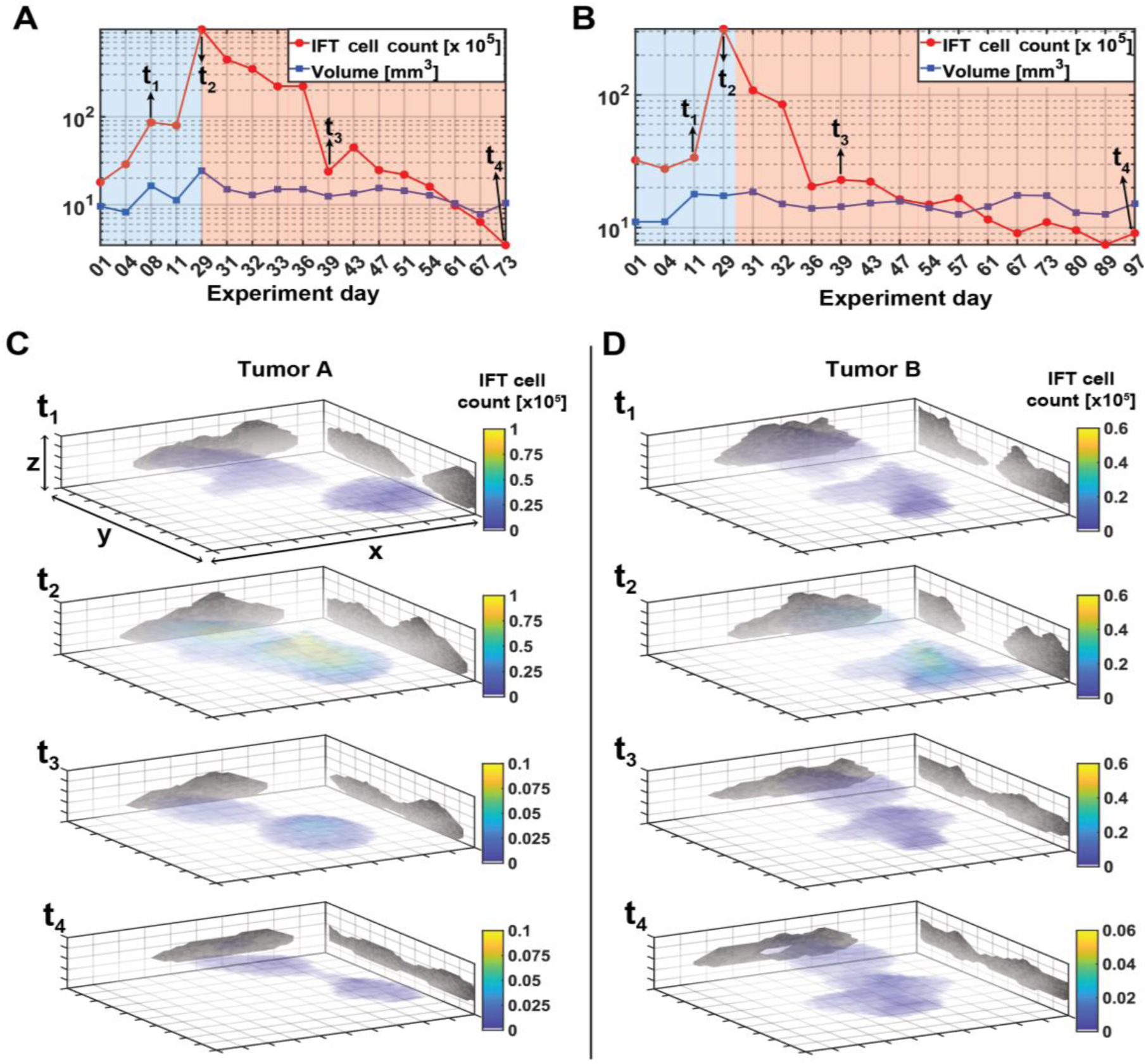
Longitudinal molecular imaging of tumor development using IFT. A.-B. depict longitudinal IFT data of cell count and tumor volume for two different animals. Blue background indicates Doxycycline administered time-frame (tumor growth), while red background indicates Doxycycline-free time-frame (tumor regression). C.-D. 3D IFT reconstructions corresponding to selected time-points. All tumors were reconstructed over the same volume, (x,y,z = 2×11×11mm3). Color bars represent the IFT cell count per voxel.

## Discussion

In this work we have presented a new, non-invasive and quantitative approach to image mammalian tumor development with molecular contrast at high mesoscopic resolution and sensitivity. Our intravital fluorescence tomography system uses a non-surgical and re-useable imaging window and thus permits prolonged observations of tumor development over several months and with arbitrary frequency, and is thus able to capture in principle the transition from tumor induction to tumor progression all the way to tumor regression upon treatment. One advantage of our method is the capability to resolve in 3D relatively small tumor cell masses comprising only few hundreds of µm and ∼10^4^ cells overall, which would normally be missed by palpation or other established methods such as ultrasound and MRI^27,49^. From a biological point of view, these capabilities could enable a close evaluation of nascent stages of tumor induction, which could be detected and 3D-reconstructed upon IFT recording. Depending on the 3D tumor geometry changes and the observed growth rate between measurements, tumors can be isolated for culture and analysis at informed, early time points that are normally not accessible for non-palpable tumors.

Similarly, the sensitivity of the IFT measurements could aid to examine stages of tumor regression, opening the possibility to capture treatment of refractory tumor clones that are not palpable anymore (e.g. time point t_3_ in **Fig. 4**). These normally inaccessible cell populations, also termed minimal residual disease, are known to be responsible for incurable tumor recurrences^4^. Hence, a handle to follow these cells and more resistant clones during the process of tumor treatment and the chance to isolate and study them in detail *in-vitro* would yield further insights on late stage tumor biology^5,8^.

Furthermore, a particular strength of our IFT technique is the ability to infer quantitative information about the underlying amount of fluorescently labeled (i.e. oncogenic) cells at relatively high accuracy. This capability sets our approach apart from other non-invasive imaging methods such as MRI, ultrasound, CT, BLI or photoacoustics, which can provide only information about overall tumor volume at mesoscopic resolution. The detailed information on the 3D geometry of tumors early in progression and during unpalpable stages of tumor regression will enable assessments of invasive or satellite states of the tumor and can be correlated with in depth histological and functional analysis upon informed resection from the animal (**Fig. 3d**).

One unexpected result of our study was the fact that tumor volume and (oncogenic) cell quantity are not well correlated – this indeed highlights the need for more quantitative approaches. A critical step in rendering IFT quantitative is the calibration to histological ground-truth data, which requires additional animals and sophisticated, time-consuming analysis. However, once calibrated, the conversion factor is expected to remain reasonably constant, and can thus be used across animals and experiments, provided that the scattering properties of the tumor and surrounding tissue and the expression of molecular fluorescence reporters does not change significantly between the calibration and experimental batches. Here, our recent work^50^ suggests that fluorophore expression and thus brightness of individual oncogenic cells stays constant over time, and is not affected by the presence of DoX. Thus, an increase in overall IFT fluorescence can indeed be attributed to an increase of oncogenic cells within a given volume. Other experimental uncertainties entails cell counting variabilities due to difference in histological staining as well as spatial inhomogeneity in tumor cell sizes and thus densities between the analyzed and extrapolated histological areas used for calibration (c.f. **Fig. 3a,b**). We estimate the overall uncertainty in absolute cell numbers to be on the order of ∼20-30%.

Unlike imaging techniques that achieve higher spatial resolution and molecular sensitivity such as confocal or multiphoton microscopy, IFT does not require surgical intervention to gain optical access to the tumor, and is capable of imaging at a much greater tissue or tumor depth. In contrast to BLI approaches, IFT does not necessitate external substrate injection during the imaging process, and thus IFT can collect data with high consistency for months without compromising the animal well-being. Combined with the relative simplicity and low-cost of the experimental apparatus, we expect IFT to become an attractive method for tracking *in-vivo* tumor progression in deep mammalian tissues in a non-invasive and longitudinal fashion. In the future, the fact that IFT can in principle record fluorescence at multiple wavelengths quasi-simultaneously should enable to study the spatial and temporal dynamics of different clonal tumor populations, their interactions as well as their effect on overall tumor burden. In addition, further functional reporters, for instance indicating apoptotic onset at a different wavelength^51^, could be employed to extract additional mechanistic information over the tumorigenesis process.

Taken together, our work combines the strengths of pre-clinical mouse models with the non-invasiveness, high resolution and sensitivity of light-based imaging methods, and thus might enable future studies on critical early stages of tumor development as well as on treatment refractory tumor clones.

## Author contributions

M.S.O., M.J. and R.P. designed research, M.S.O., M.G.M., L.W and L.M.C. performed research, M.S.O., wrote software, M.S.O., and M.G.M. analyzed data; M.S.O., M.J. and R.P. wrote the paper. M.J. and R.P. supervised research.

## Data and code availability

The datasets generated and software code utilized in the current study are available from the corresponding author on reasonable request.

## Acknowledgments

This work was supported by the European Molecular Biology Laboratory (EMBL), a fellowship to M.S.O. under Marie Skłodowska-Curie Actions COFUND (grant agreement number 664726). We would further like to thank the EMBL mouse facility and mechanical workshop; M. Paulsen from the Flow Cytometry Core Facility, as well as the group of R. Sotillo (DKFZ) for their support on tumor slice imaging.

## Methods

### Transgenic mouse model

Mice used in this study were bred and maintained in the EMBL-Heidelberg animal facility (IACUC protocol MJ160070), in accordance to the guidelines of the European Commission, revised Directive 2010/63/EU and AVMA Guidelines 2007.

Mouse strains utilized in this study: RAG1 (-/-), as immunodeficient mouse receiver strain ^52^ (C57BL/6 background), and the trackable tumor donor strain TetO-cMYC/TetO-Neu/MMTV-rtTA/H2B-mCherry [FVB background] (i.e. tumorigenic cell nuclei are labelled with the fluorescence protein mCherry). In order to generate the donor strain, the tri-transgenic mouse line TetO-cMYC/TetO-Neu/MMTV-rtTA^6^ was bred with the R26-H2B-mCherry strain^53^.

All efforts were made to minimize the amount of animals used - in total 30 mice for the experiments lined out in this manuscript - in accordance with Russell and Burch’s principle of (3Rs) reduction and highest ethical standard. Animals were kept on a 12-hour light/12-hour dark cycle, with constant ambient temperature (23±1°C) and humidity (60±8%), supplied with food pellets (for tumor induction, pellets contained Doxycycline hyclate, 625 mg/kg; Envigo Teklad) and water ad libitum.

### Cells transplantation

Mammary glands from the donor strain (female virgin mice 7-9 weeks old) were harvested, singularized and seeded on a 2D collagen coated 6-well plate (BD, Cat. # 356400). Cells were fed with mammary gland specific media (Promocell, Cat.#21210), plus Doxycycline hyclate (Sigma, D9891) overnight. The next day, tumor cells were extracted, counted and resuspended in a solution of PBS and Matrigel (Corning, 356231). A mixture of 150.000 cells in 40µl of media were injected into the donor strain using the 50µl syringe (Hamilton, 705N ga22s/51mm/pst2). Female virgin RAG1 animals (from 4-8 weeks), were fed with Doxycycline diet (Envigo, TD.01306) at least two days prior to transplantation and for as long as tumor growth was desired. Animals were anesthetized using Xylazine/Ketamine before the surgery procedure. Closure of the wound was done by manual suture and no further post operational issues or inflammatory responses were observed.

### Intravital fluorescence tomography

In order to reconstruct a 3D image from the raw 2D data, IFT proceeds as follows. First, a so-called forward model (Jacobian matrix) is calculated, in which photon propagation inside the scattering tissue is simulated via a GPU-accelerated Monte-Carlo method^37,44^ <u>(MCX, 2019.4</u>, GeForce GTX 1080, 8MB) (see **Fig. 1b,iii**). The Jacobian matrix was computed with 10^8^ photons and a 0.25 × 0.25 × 0.1 mm^3^ voxel size. Computation took ∼3 minutes on a desktop PC (Intel Core i7-6800K @ 3.4 GHz). The total simulation volume for the *in-vivo* experiments was 10×10×2mm^3^). The integrating sphere experiments **(Supplementary Note 1 and Supplementary Fig. S1 & S2)** showed that any differences in optical properties between the excitation (561nm) and maximum emission wavelength (610nm) is less than 5%. Therefore, we used the same optical property values for the Jacobian simulations. For the majority of the results presented here, the following parameters were used: μ^s^ = 20mm^−1^, µ^a^=0.1mm^-1^, g=0.85. This ensured accurate and reliable 3D reconstructions. For solving the inverse problem of the image reconstruction, the following cost function was used, which delivered an optimized solution for the target volume *x*:

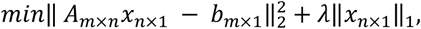

Here *A* is the Jacobian matrix and *b* the experimental imaging dataset. The regularization parameter, *λ*, is a penalizing term which acts as a filter for *x* to prevent noise from dominating the reconstruction. An initial value for *λ* was found by utilizing the L-curve method^54^. Then, a fine tuning was conducted based on the similarity coefficient^55^ between the raw data and the maximum intensity projection of the reconstruction. The regularization parameter (*λ*) was chosen based on the highest similarity value. This procedure allowed us to account for slight differences in noise levels between individual datasets, while minimizing any user bias in the reconstruction. Before 3D reconstruction, the collected raw images were sub-divided into smaller images depending on the desired source-detector (S-D) separation, which is proportional to the targeted imaging depth and inversely proportional to both lateral and axial resolution. In our work, the chip size of the sCMOS camera allowed us to place detectors up to 3.5mm away from the source (aligned to the center of the camera). With these parameters, the system is capable of reconstructing deeply seated fluorescent emitters from up to 3mm below the surface.

### Optical setup

A schematic overview of the experimental arrangement is given in **Fig. 1c**. An excitation laser (561nm) was coupled into the setup by a dual-band dichroic mirror (Di488/561, Semrock). A 2D galvanometric mirror pair (GVS012/M, Thorlabs), scans the excitation spot on the sample via a scanning lens (2x AC508-200, f=200mm, 2”, in Plössl configuration). The Plössl design and large aperture optics in our setup yield a particularly large scanning area and thus FOV, i.e. up to 15×15mm^2^ (diameter = 22mm) with a 1x effective magnification. The emitted fluorescence light is descanned by the same 2D mirror and imaged on the detector array (sCMOS, Zyla 4.2 Plus, Andor) after passing a dual-band emission filter (FF01-523/610-25, Semrock). For most experiments in this work, we used 512×512 active pixels on the camera. The high speed of the sCMOS camera enabled fast data collection (∼16 seconds for a grid of 41×41 scan points and thus 1681 images, ∼881MB). Laser power focused on the sample ranged between 1-5mW with 10ms dwell time for each scan point. The energy density of the laser was kept between 15-150mJ/cm^2^ which even in the worst case is ∼9 times less than the established Maximum Permissible Exposure level (1.3 J/cm^2^) according to the standards (60825-1/A2 standard in Appendix 1) for the skin.

### IFT imaging procedure

Mice were imaged according to approved EMBL IACUC protocols. Each animal was anesthetized for the duration of the imaging via Isoflurane. Once heartbeat and breathing stabilized, a topical hair removal cream was applied covering an area of ∼5cm^2^ revealing the abdominal mammary gland. The imaging procedure consisted of an initial quick scan over the entire area of interest (4th and 5th mammary gland) to identify locations of fluorescence and thus potential tumor initiation. Once identified, we applied the non-invasive imaging window onto the area and acquired the raw IFT data. Typically, this entire process took ∼15-20min, of which the raw data acquisition time only took <20sec. During the initial stages of tumor induction, imaging was performed weekly with more regular monitoring, up to daily, during the last days of growth and first days of regression phases. During the late phases of tumor regression, we again reduced the imaging frequency to weekly or bi-weekly intervals.

### Histology and H&E processing

The H&E staining refers to the mixture of Hematoxylin (Vector, H-3404) and Eosin Y (Fisher, LAMB/100-D), for nuclear and cytoplasm staining, respectively. Tissue histology and H&E staining were performed as outlined below, and following Ref.^56^. Tumors were extracted and fixed using 4% formalin (Sigma, HT501128), for 24 hours at room temperature, followed by exposure to increasing concentrations of ethanol up until 100%, after which the sample is moved to xylene for clearing and finally embedded in paraffin. The paraffin block was then sectioned using a microtome to obtain single, 4.5µm thick slices. Slides containing section slices were cleaned from paraffin using xylene (Roth, CN80.2), and then hydrated using a decreasing concentration of ethanol. The samples were exposed to Hematoxylin for 90 secs and to Eosin for 30 secs. After going through washes, the slides were once more put in contact with an ascending concentration of ethanol, ending on xylene. Slides were mounted using DPX mounting media (VWR, 360294H). As a consequence of the heterogeneous nature of the tumor phenotype, this staining protocol yielded stainings of variable outcome. In particular, only a small percentage of tumors had well-stained nuclei, which are a prerequisite to separate them from the cytoplasm and thus ensure accurate counting with commercial image processing software (HistoQuest software, Tissuegnostics, Vienna, Austria). This motivated us to devise a custom image processing pipeline (see **Fig. 3a,b**).

## Supplementary Information

### Supplementary Note 1: Double-integrating-sphere setup for measuring bulk optical properties of mouse skin tissue

To measure bulk optical properties of mouse tissue, we utilized a double-integrating sphere setup (DIS) with the sample in the middle ^1,2^ (Error! Reference source not found.). From transmittance, diffuse and reflected light measurements the optical properties can be estimated using the inverse adding doubling (IAD) method proposed by Prahl et. al. ^3,4^. Each integrating sphere (EverFine Corporation, Hangzhou, China) was equipped with a ϕ=3.6 mm Si photodiode (PDA36A2, Thorlabs Inc., New Jersey, USA) whose voltage readings were digitized by a 16-bit DAQ card (PCIe-6323, National Instruments, Texas, US). The inner walls of the integrating spheres were coated with barium sulfate and had a diameter of 30 cm. A supercontinuum laser (SC-Pro-7, YSL Photonics, Wuhan, China) in combination with an acoustic-optic tunable filter (AOTF) device (AOTFnC-VIS, AA Opto-Electronics, Orsay, France) provided a rapid wavelength scanning light source over the band of 550 – 720 nm. The power spectra (Error! Reference source not found. inset) at the fiber exit port were calibrated by combining a spectrometer (USB4000, Ocean Insight, US) and a powermeter (PM100D, Thorlabs Inc., New Jersey, USA). Immediately after the AOTF, the monochromatic light beam was coupled into a 50 μm core diameter multi-mode fiber. A collimator lens with f = 11 mm generated a round light spot (ϕ ∼ 3 mm) on the sample positioned between the spheres. Detector ports are masked from the sample and entrance ports by internal baffles, in order to ensure the detection of diffuse light only. Because of the sphere geometry, the sample is illuminated under an angle of 9°. Both diffuse and reflected (including specular reflectance) light are measured together by the detectors. We used a custom-designed and 3D-printed sample holder made from black polyether ether ketone (PEEK) which fit the sphere ports precisely and had a ϕ = 10 mm access hole. Before measurements, the light beam was aligned to be focused at the center of the access hole. The skin sample was taken from the mammary gland area of the sacrificed mouse (following FELASA guidelines). First, the mouse skin was separated from the peritoneum, avoiding any tissue shrinkage and then was sandwiched by two microscope glass slides with thickness of 1.00 mm. Once a sample was inserted into the sample holder and positioned between the spheres, measurements were performed by home-made LabView control program which synchronized the digitalization to the wavelength scanning. Data for each wavelength were averaged from 1,000 consecutive measurements with a 10kHz sampling rate. An entire measurement over the whole wavelength range could be finished within 1 minute. The measurements along with the calibration data were preprocessed followed by the optical property estimation using the IAD approach. Error! Reference source not found. (a) and (b) show the so measured scattering and absorption coefficients of two mouse skin samples, respectively.

**Supplementary Figure 1:**
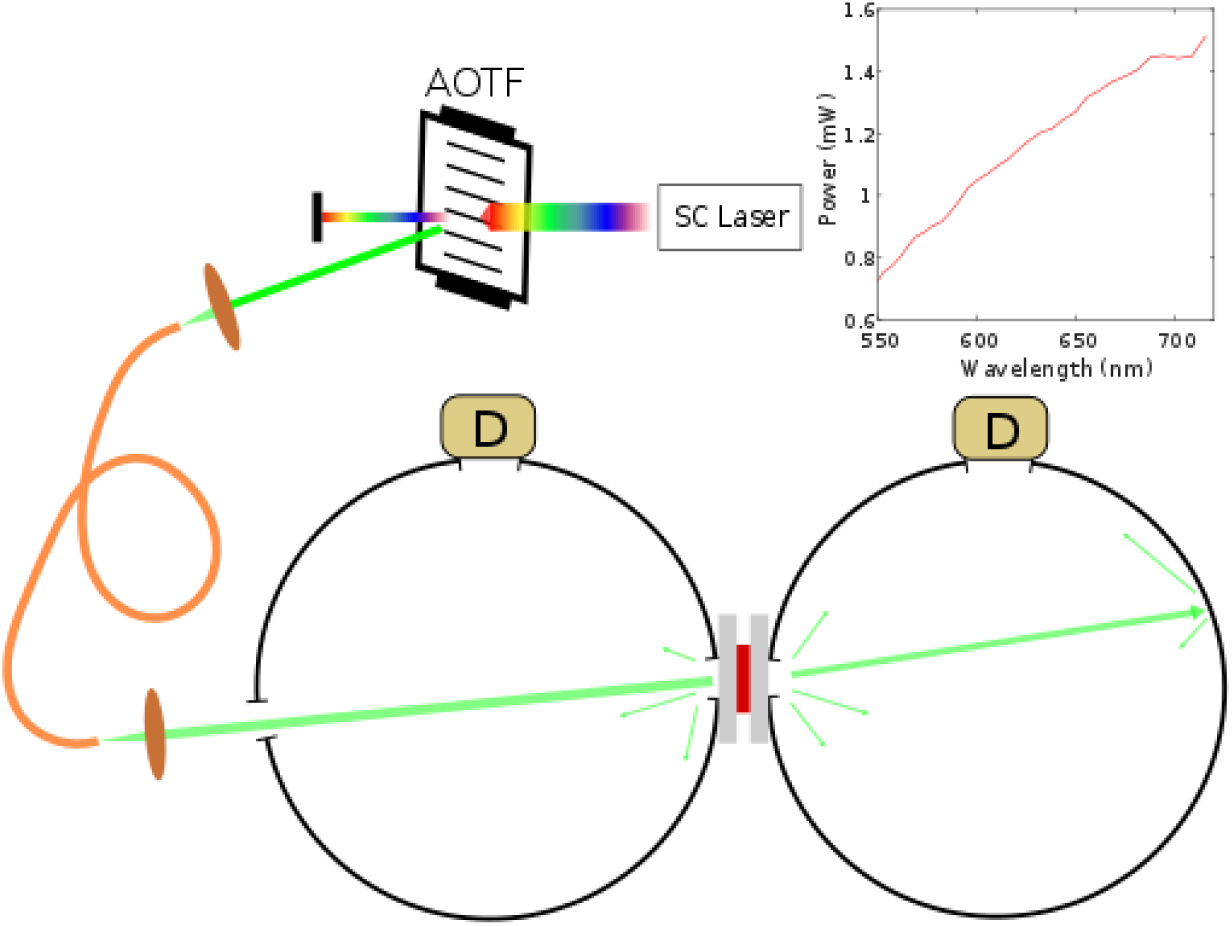
Schematic overview of the double integrating sphere system in the 550-720 nm range. Inset: Light power spectrum at the fiber exit port when the SC laser functions at the maximum output. SC laser = Supercontinuum laser; D = photodiode detector; AOTF = Acoustic optic tunable filter.

**Supplementary Figure 2:**
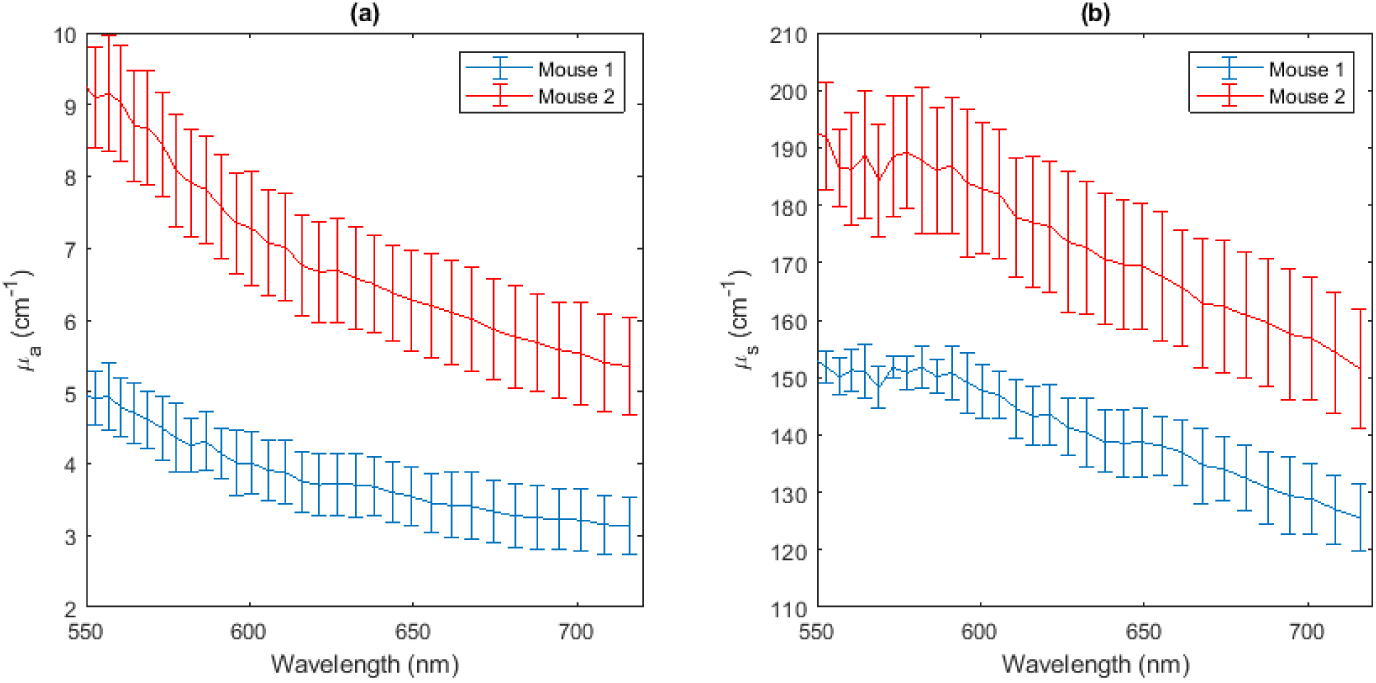
Optical properties of mouse skin. (a) Absorption coefficient and (b) scattering coefficient in the 550-720 nm range. Data are averaged from 3 measurements of different locations and the error bars denote standard deviation. The skin thicknesses of mouse 1 and 2 were 0.55 mm and 0.50 mm, respectively, as measured by a caliper. The anisotropy factor (g) and refractive index of skin were assumed as 0.9 and 1.4, respectively ^5^.

### Supplementary Note 2: OCT imaging of mouse skin

The performance of the NIVOW was evaluated by a self-built spectral-domain Optical Coherence Tomography (SD-OCT) setup with a linear wavenumber space spectrometer, whose design follows reference ^6^. Briefly, the spectrometer of the SD-OCT setup utilized a 28 kHz line camera (AViiVA SM2 CL, e2v, Cedex, France) covering a wavenumber range of 8.267–6.830 μm^−1^ (center wavelength 832 nm). The measured axial resolution in tissue (n=1.35) was ∼3.5 μm. The transversal light beam scanning was achieved by a combination of galvo mirror pair and a telecentric scanning lens (f= 60.3 mm). When the laser beam diameter was 4.0 mm, the transversal resolution at the focal plane was measured to be 23.8 μm. Intensity profile of depth scan (A-line) was reconstructed from interference spectral signal following a regular OCT postprocessing procedure including steps of dispersion compensation, background subtraction, spectrum reshaping and inverse fast Fourier transformation ^7^. Each cross-sectional image (B-scan) contained 128 A-lines of 2048 depth samples, and took ∼4.57 ms to acquire. The time-lapse images were acquired with a rate of 54.8 frames/s. The transversal spatial scale of the OCT imaging system was calibrated by placing an array camera (CM3-U3-50S5M-CS, FLIR, Wilsonville, USA) at the focal spot while the axial scale was calibrated by a 1.00 mm thickness air gap between two regular microscope slides. In order to quantify the tissue movement during the experiment, a cross section of the mammary gland tissue area was imaged by OCT at full speed (**Supplementary Video 1 & 2**). From these movies every consecutive image pair of the image series was digitally registered with a resolution of 1/10 pixel followed by pixel-micrometer conversion with the calibrated spatial scale factors in both axial and transversal directions ^8,9^. The spatial shifts of each frame relative to the first were then plotted in Fig. 1E.

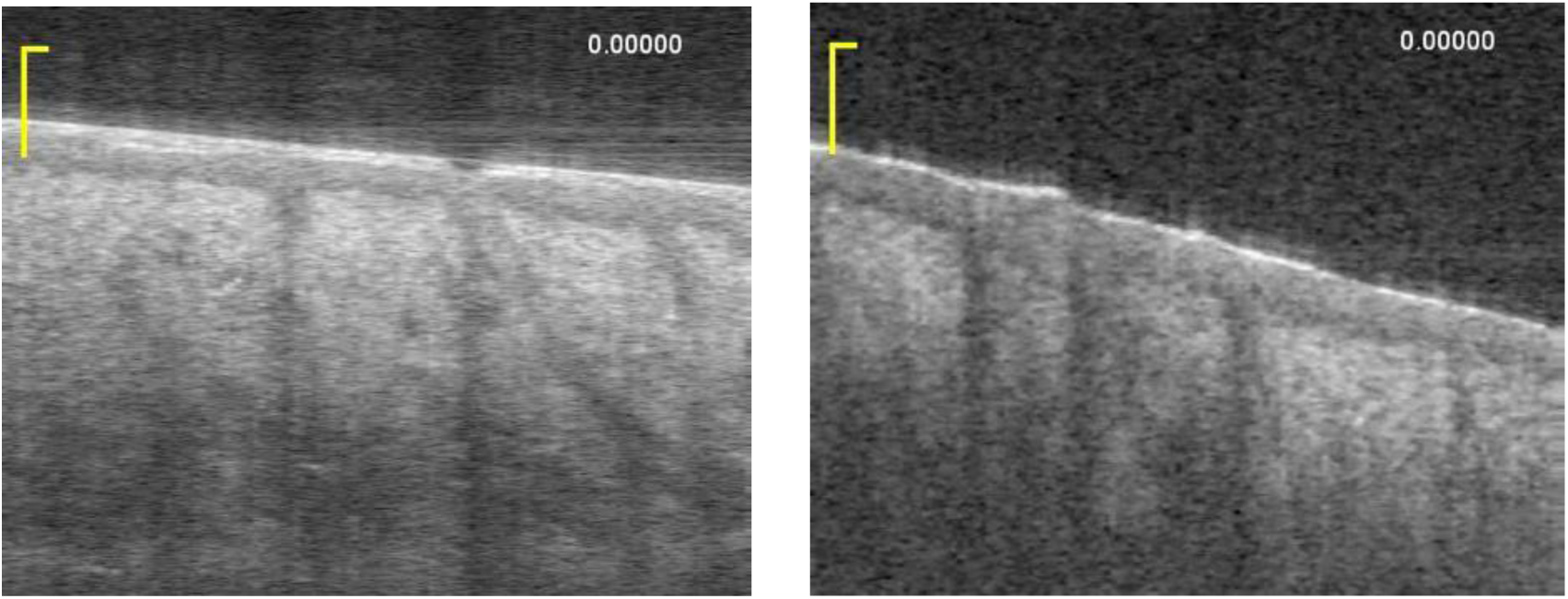
Time-lapse cross-sectional images of mouse abdomen skin over 9.4 s obtained by the OCT setup without (left) and with (right) the NIVOW. Scale bar 100 μm. See **Supplementary Videos 1 & 2**.

